# Regional low-frequency oscillations in human rapid-eye movement sleep

**DOI:** 10.1101/397224

**Authors:** Giulio Bernardi, Monica Betta, Emiliano Ricciardi, Pietro Pietrini, Giulio Tononi, Francesca Siclari

**Author notes:** Correspondence: Francesca Siclari, MD, CHUV, Centre d’investigation et de recherche sur le sommeil, Rue du Bugnon 46, CH-1011, Lausanne, Switzerland,; Giulio Bernardi, MD, PhD, IMT School for Advanced Studies Lucca, Piazza S.Francesco, 19 IT-55100, Lucca, Italy.

## Abstract

Although the EEG slow wave of sleep is typically considered to be a hallmark of Non Rapid Eye Movement (NREM) sleep, recent work in mice has shown that slow waves can also occur in REM sleep. Here we investigated the presence and cortical distribution of low-frequency (1-4 Hz) oscillations in human REM sleep by analyzing high-density EEG sleep recordings obtained in 28 healthy subjects. We identified two clusters of low-frequency oscillations with distinctive properties: 1) a fronto-central cluster characterized by ∼2.5-3.0 Hz, relatively large, notched delta waves (so-called ‘sawtooth waves’) that tended to occur in bursts, were associated with increased gamma activity and rapid eye movements, and upon source modeling, displayed an occipito-temporal and a fronto-central component; and 2) a medial occipital cluster characterized by more isolated, slower (<2 Hz) and smaller waves that were not associated with rapid eye movements, displayed a negative correlation with gamma activity and were also found in NREM sleep. Thus, low-frequency oscillations are an integral part of REM sleep in humans, and the two identified subtypes (sawtooth and medial-occipital slow waves) may reflect distinct generation mechanisms and functional roles. Sawtooth waves, which are exclusive to REM sleep, share many characteristics with ponto-geniculo-occipital (PGO) waves described in animals and may represent the human equivalent or a closely related event while medio-occipital slow waves appear similar to NREM sleep slow waves.

## Introduction

Sleep is characterized by relative quiescence and reduced responsiveness to external stimuli. Based on electrophysiological hallmarks, sleep is divided into non-rapid eye movement (NREM-)sleep and REM-sleep. While REM-sleep is characterized by rapid eye movements and muscular atonia, and by a tonically ‘activated’ (low voltage, high frequency) EEG resembling that of wakefulness [1,2], hallmarks of NREM-sleep include high-amplitude, slow waves (≤4 Hz) and spindles (12-16 Hz) [3,4]. Given these differences, it has been commonly assumed that wakefulness, NREM-sleep and REM-sleep represent mutually exclusive ‘global’ states. This view has been recently challenged by a growing body of evidence indicating that many features of sleep are essentially local, and that islands of sleep- and wake-like activity may coexist in different brain areas [5].

EEG slow waves of NREM-sleep occur when neurons become *bistable* and oscillate between two states: a hyperpolarized *down-state* characterized by neuronal silence (*off-period*), and a depolarized *up-state* during which neurons fire (*on-period*) [4]. Intracranial recordings in humans have shown that during stable NREM-sleep, slow waves can be restricted to some localized areas and occur out of phase with other cortical regions [6]. The regional distribution of slow wave activity (SWA) has been shown to be modulated by recent experience and learning (e.g., [7,8]). Local slow waves, involving small portions of the cortical mantle, have also been shown to occur in awake humans [9,10] and rodents [11,12], particularly in conditions of sleep deprivation. Importantly for the purpose of our study, recent work demonstrated that in mice [13], slow waves with neuronal *off-periods* may also occur during REM-sleep in primary visual (V1), sensory (S1) and motor (M1) areas, mainly in cortical layer 4. By contrast, associative areas, such as V2, S2, or the retrosplenial cortex, showed the typical activated pattern of REM-sleep. It is currently unknown whether similar regional differences and local slow waves also exist in human REM-sleep. In fact, while great effort has been dedicated to the study of prominent features of REM-sleep, such as theta (5-8 Hz) or high-frequency (>25 Hz) activity, little attention has been given to low-frequency (≤4 Hz) oscillations during this sleep stage. A few studies found that *delta* power (1-4 Hz) in REM-sleep had a different topographic distribution with respect to NREM-sleep [14,15]. In addition, we recently showed that both in REM and NREM sleep, reduced *delta* power in posterior cortical regions is associated with dreaming [16], suggesting a similar functional significance of slow waves across states. By analyzing slow wave characteristics in NREM sleep, we identified two types of slow waves with distinct features: large, steep fronto-central type I slow waves, which are likely generated in a bottom-up manner by arousal systems, and smaller type II slow waves, which are diffusely distributed over the cortical mantel and probably underlie a cortico-cortical synchronization mechanism [17]. Further analyses showed that the two types of slow waves are differentially related to dreaming [18].

In the present study we aimed at investigating the presence and cortical distribution of low-frequency (LF; 1-4 Hz) oscillations in human REM-sleep using high-density EEG recordings, which provide an excellent combination of temporal and spatial resolution. We also wanted to determine whether we could identify different LF wave types with distinct characteristics, similarly to NREM sleep. Sawtooth waves of REM-sleep, for instance, are known to peak in the *delta* range (∼3 Hz), but their properties and cortical distributions have not been systematically investigated in humans using techniques with a high spatial resolution. Therefore, we expected to identify at least two types of LF oscillations in human REM-sleep, corresponding to sawtooth waves and local slow waves involving primary sensory cortices.

## Material and Methods

### Participants

Overnight high-density (hd-)EEG recordings (Electrical Geodesics, 257 electrodes, 500 Hz sampling frequency) were obtained in 16 healthy volunteers (age 24.9±4.5, 8 females; 13 right-handed). NREM-sleep data (but not REM-sleep data) from 10 of these subjects has been reported in a previous publication [19]. All participants had a sleep duration of ∼7 h/night, consistent bed/rise times, no daytime nap habits and no excessive daytime sleepiness (total scores in the Epworth Sleepiness Scale ≤10; [20]). A clinical interview was performed to exclude a history of sleep, medical and psychiatric disorders. The study was approved by the IRB of University of Wisconsin-Madison. Each participant signed an IRB-approved informed consent form before enrollment into the study.

Since wake data was not available from the above subjects, specific analyses aimed at comparing sleep- and wake-related brain activity (see Methods) were performed in a different sample of 12 healthy adult individuals (age 25.5 ± 3.7, 6 females) who underwent overnight hd-EEG sleep recordings after having spent 8 h in the sleep laboratory while watching movies (analyses of wake and NREM-sleep data from these subjects has been reported in [21]). These recordings were performed at Lausanne University Hospital, under a research protocol approved by the local ethical committee.

### EEG data preprocessing

All hd-EEG recordings (Madison dataset) were first-order high-pass filtered at 0.1 Hz and band-pass filtered between 0.5 and 58 Hz. For scoring purposes, four of the 257 electrodes placed at the outer canthi of the eyes were used to monitor eye movements (electrooculography, EOG), while electrodes located in the chin-cheek region were used to evaluate muscular activity (electromyography, EMG). Sleep scoring (Table 1) was performed over 30s epochs according to standard criteria by a sleep medicine board certified physician [22]. All the epochs scored as REM-sleep were identified and extracted.

**Table 1.** Sleep parameters (average and standard deviation) for the 16 subjects included in the study.

| Parameter (N=16) | Average | SD |
| --- | --- | --- |
| Total Sleep Time (min) | 397.5 | 79.8 |
| N1 Time (min) | 21.8 | 15.2 |
| N1 Proportion (%) | 5.6 | 4.2 |
| N2 Time (min) | 209.5 | 52.1 |
| N2 Proportion (%) | 52.9 | 9.0 |
| N3 Time (min) | 78.0 | 29.6 |
| N3 Proportion (%) | 20.0 | 6.5 |
| REM Time (min) | 88.3 | 33.3 |
| REM Proportion (%) | 21.5 | 6.1 |
| REM Latency (min) | 91.5 | 33.5 |
| REM Cycles (n) | 4.0 | 1.4 |

### Detection and classification of low-frequency oscillations

For each subject and recording, channels containing artifactual activity were visually identified, rejected, and replaced with data interpolated from nearby channels using spherical splines (NetStation, Electrical Geodesics Inc.). Potential ocular, muscular and electrocardiograph artifacts were removed using Independent Component Analysis (ICA) in EEGLAB [23]. An algorithm based on the analysis of consecutive signal zero-crossings [24] was adapted for the detection of low-frequency oscillations in REM-sleep. Before application of the algorithm, the signal of each channel was referenced to the average of the two mastoid electrodes and filtered between 1 and 10 Hz (the upper limit of the filter was selected to minimize wave shape and amplitude distortions). For each detected half-wave with duration between 125 and 500 ms (1-4 Hz), the following properties were extracted and stored for further analyses: negative amplitude (µV), duration (ms) and number of negative peaks (np/w). The density of slow waves (waves per minute; w/min) was also computed.

Preliminary analyses revealed two spatially distinct clusters of LF waves in the 1-4 Hz range: a fronto-central and a medio-occipital cluster characterized by different frequencies and amplitudes (Figure 1). For this reason, subsequent analyses were in part adapted to the characteristics of the two LF wave types (see Results).

**Figure 1.**
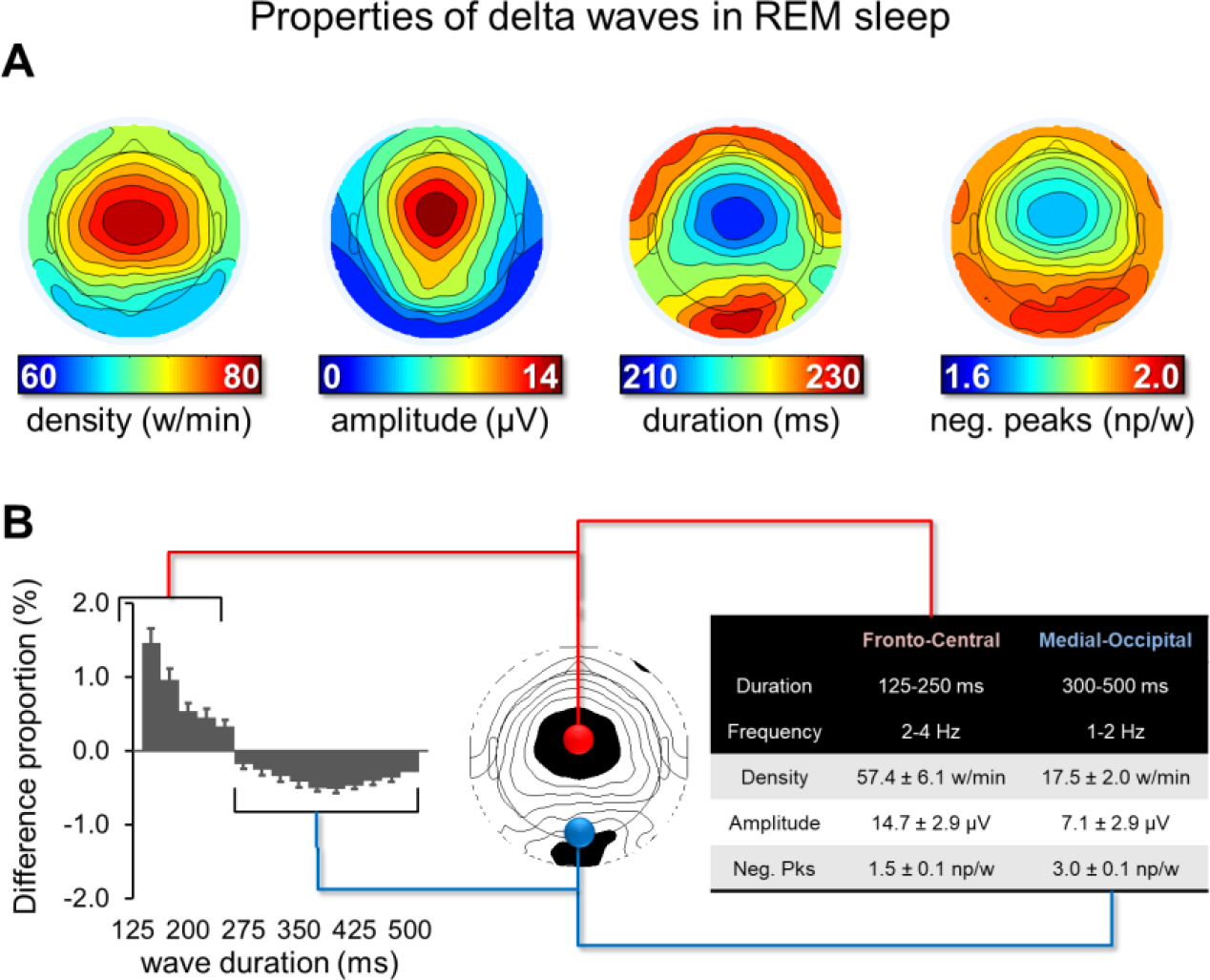
Properties of LF waves in REM-sleep. An automatic detection algorithm was applied to the EEG signal of each electrode to identify negative half-waves with a duration of 125-500 ms (≤4 Hz). Different wave properties were extracted and analyzed (panel A), including density (number of waves per minute; w/min), negative amplitude (µV), duration (ms) and number of negative peaks (np/w). Panel B (left) shows, for a fronto-central and a medial-occipital electrode, the difference (%) in the relative proportion of waves for durations between 125 and 500 ms (25 ms bins). The relative difference between the two electrodes in % was calculated for each duration bin. Vertical bars indicate one standard deviation from the group mean (SD). The table on the right side displays mean values and SD for density, amplitude and negative peaks (neg. pks) for the faster fronto-central and the slower medio-occipital LF waves.

### Temporal clusterization of low-frequency oscillations

We investigated the tendency of LF waves to occur in bursts. Bursts were defined as a series of at least 3 consecutive waves whose maximum negative peaks were less than 750 ms apart. For this analysis a slightly broader half-wave duration limit (range 1-5 Hz) was used to minimize exclusion of potential ‘borderline’ LF waves within a burst. An amplitude constraint was applied so that the amplitude of each consecutive wave in the burst had to vary by less than 25% (relative to the preceding wave). No other amplitude or duration threshold was applied.

### Relationship with rapid eye movements

Individual rapid eye movements (EMs) were identified using an automatic detection algorithm [25,26]. In brief, the algorithm detects local peaks in the sum of the Haar Wavelet Transform coefficients in a specific frequency range (0.8-5 Hz) to determine the exact timing and duration of each eye movement [25]. Bursts of rapid EMs were defined as serial EMs (2 or more) separated by less than 1 s. *Isolated* rapid EMs/bursts were defined as single EMs/bursts preceded and followed by periods of at least 5s without rapid EMs. To determine how LF waves relate to rapid EMs, for each isolated EM, wave density was calculated in two windows corresponding to 1 s before the onset of the first EM (‘pre’) and 1 s after the end of the last EM (‘post’).

Phasic REM periods were defined as time periods longer than 5 s that were occupied by EMs for more than 50% of their length. EMs in this time window had to be less than 2s apart to be considered as part of the same phasic period. Tonic REM episodes were defined as time periods longer than 5s that were completely free from any EMs. The average density of fronto-central and medial-occipital waves was calculated in all phasic and tonic periods.

In order to increase the signal-to-noise ratio of these analyses, amplitude thresholds corresponding to 10 µV and 3 µV were applied for fronto-central and medial-occipital LF waves, respectively.

### Relationship with high-frequency activity

Next, in light of previous observations linking NREM slow waves to changes in high-frequency EEG activity [17,27], we examined the temporal relation between *gamma* and *delta* power during LF waves of REM-sleep. For each wave, the band-specific instantaneous signal power was computed as the root mean square (RMS, time-window = 40 ms, step = 20 ms) of the *delta* (0.5-4 Hz) and *gamma* pass-band filtered (30-55 Hz) signals for the 2 s before and after the negative peak of the LF wave. To increase the signal-to-noise ratio, amplitude thresholds corresponding to 10 µV and 3 µV were applied for fronto-central and medial-occipital LF waves, respectively. For each subject, *delta* and *gamma* RMS time-series were z-score transformed and averaged across all the detected waves. A group average was computed on the 600 ms time window centered on the negative peak of the LF wave.

### Overnight changes in low-frequency oscillations

Slow waves of NREM-sleep are known to change in density and amplitude across a night of sleep [19,24]. Thus, here we evaluated whether LF waves of REM-sleep undergo similar changes. Specifically, for each subject, we first calculated the mean density and amplitude of LF waves within each REM-sleep cycle. Then, the magnitude of the density/amplitude changes (‘slope’) was calculated across cycles using least square linear regression (considering as independent variable the absolute sleep time at half of the corresponding REM-sleep cycle).

### Relationship with low-frequency oscillations of NREM-sleep

In order to evaluate similarities and differences between LF waves of REM- and NREM-sleep, additional analyses were performed on a sub-sample of 10 subjects (4 females, age 25.4±4.7). NREM data of these 10 subjects was preprocessed using the same procedures described above as part of a previous publication aimed at exploring different types of slow waves during NREM-sleep [19]. For these subjects, the same negative half-wave detection algorithm described above was applied to N2 and N3 sleep data of the whole night on a channel-by-channel basis. Then, half-waves with duration 125-250 ms (>2 Hz) and 300-500 ms (<2 Hz) were identified to calculate mean density (w/min) and negative amplitude (µV). A non-parametric Friedman test was applied to investigate potential stage-dependent (N2/N3/REM) differences in these parameters.

### Specificity of REM-sleep low-frequency oscillations with respect to wakefulness

In order to verify whether LF waves of REM sleep are specific of this stage or may also be observed in wakefulness, additional analyses were performed using REM and waking (rest with eyes closed) data collected in 12 healthy adults (6 females, age 25.5 3.7; [21]). These recordings were preprocessed as previously described, but a 0.5-45 Hz bandpass filter was used in this case. Moreover, resting state recordings in wakefulness were initially divided into non-overlapping 5 s epochs and visually inspected to identify and reject epochs containing clear artifacts, before application of the ICA procedure. A non-parametric Wilcoxon signed rank test was used to compare amplitude and density of LF waves across REM-sleep and wakefulness.

### Source modeling analysis

A source modeling analysis was performed to determine the cortical involvement of LF waves in REM-sleep. First, the negative-going envelope of the EEG signal was obtained by selecting the second most negative sample across 65 channels included in a medial centro-frontal or in a temporo-occipital region of interest (ROI) [17,28]. The resulting signal was broadband filtered (0.5-40 Hz, stop-band at 0.1 and 60 Hz) and the same negative half-wave detection algorithm described above was applied. Specific criteria were applied to further select waves with typical properties of fronto-central and medial-occipital waves, respectively, and to optimize source estimation while minimizing the potential impact of residual noise. In particular, anterior waves with a negative amplitude greater than 20 µV and a duration of 125-375 ms, and posterior waves with a negative amplitude of 5-50 µV and a duration >300 ms were analyzed. The duration ranges applied here were broader than those reported for the other analyses at the single-channel level in order to take into account potential wave propagation [28,29]. For the analysis of fronto-central waves, the EEG signal was filtered between 2 and 5 Hz (stop-band at 1.5 and 6 Hz), while it was filtered between 0.5 and 2.5 Hz (stop-band at 0.1 and 4 Hz) for medial-occipital waves. Then, source localization was performed on 1s data segments centered on the negative peak of each LF wave using GeoSource (NetStation, Electrical Geodesics, Inc). A four-shell head model based on the Montreal Neurological Institute (MNI) atlas and a standard coregistered set of electrode positions were used to construct the forward model. The inverse matrix was computed using the standardized low-resolution brain electromagnetic tomography (sLORETA) constraint. A Tikhonov regularization procedure (10^-2^) was applied to account for the variability in the signal-to-noise ratio. The source space was restricted to 2,447 dipoles distributed over 7 mm^3^ cortical voxels. For each LF wave, the cortical involvement was computed as the average of the relative current density achieved within a time-window of 20 ms around the wave negative peak. Finally, for each subject, the probabilistic involvement was defined as the probability for each voxel to be among the 25% of voxels showing the maximal cortical involvement.

### Experimental Design and Statistical Analysis

All statistical analyses were performed using MATLAB (TheMathWorks, Inc.) or SPSS Statistics (IBM Corp., Armonk, N.Y., USA).

Statistical comparisons between phasic and tonic REM periods or between pre- and post-EM periods were performed using parametric paired t-tests including all study participants (N=16). The relationship between the number of LF waves and the number of rapid EMs was investigated using the Spearman’s correlation coefficient. A non-parametric, permutation-based test was used to evaluate significance of overnight amplitude and density changes in LF waves. Non-parametric Friedman’s tests were applied to identify potential differences across sleep stages (N2, N3, REM), while Wilcoxon signed rank tests were used for paired comparisons with relatively small sample sizes (N=10, or N=12).

When hypotheses were tested across multiple voxels/electrodes, a non-parametric approach based on the definition of a cluster-size-threshold was used to control for multiple comparisons [30,31]. Specifically, a null distribution was generated by randomly relabeling the condition-label from the original data for group-level comparisons and by randomly reordering observations for correlation analyses (N permutations = 1,000). For each permutation, the maximal size of the resulting clusters reaching an (uncorrected) significance threshold of p < 0.05 were included in the cluster-size distribution. Then, the 95th percentile (5% significance level) was determined as the critical cluster-size threshold.

## Results

### Topographic analysis of low-frequency oscillations in REM-sleep

First, we evaluated the topographic distribution of the density, amplitude, duration and negative peaks of all negative waves detected in the 1-4 Hz range (Figure 1A). We identified two spatially distinct clusters with different properties: i) a fronto-central cluster, in which waves had a relatively high density, a high amplitude, a low duration and a low number of negative peaks, and ii) a medial-occipital cluster characterized by fewer waves, with a lower amplitude, a longer duration and a higher number of negative peaks. Next, we selected a representative electrode for each of the two clusters (channel 8, corresponding to the channel between Fz and Cz, and channel 137, corresponding to Oz), and compared the distribution of wave density as a function of their duration between the two channels (Figure 1B, left part). This analysis confirmed that the frontal cluster contained more faster waves (125 to 250 ms, >2 Hz), while the medial-occipital cluster contained more slower waves (300 to 500 ms, <2 Hz). Mean values for density, amplitude and negative peaks for characteristic faster fronto-central waves (125-250 ms) and slower medio-occipital waves (300 to 500 ms) are shown in Figure 1B (right part). Next, to better characterize the temporal dynamics of the occurrence of LF waves, we evaluated the topographic distribution of wave parameters only for LF waves occurring in bursts (see Methods for details). This analysis revealed that waves occurring in bursts were mainly found in the fronto-central cluster and had similar characteristics to the fronto-central waves described above (Figure 2). For subsequent analyses, we distinguished between fronto-central faster waves lasting 125-250 ms, and medio-occipital slower waves lasting 300-500 ms. We found that both types of waves were essentially local, as indicated by the mean proportion of recruited channels (32.0 ± 4.0% for fronto-central waves and 9.5 ± 3.5% for medial-occipital waves; Figure 3). Importantly, faster fronto-central and slower medial-occipital waves tended to occur independently from each other: co-occurrence of oscillations in a fronto-central and a medial-occipital electrode was observed for only 15.2 ± 5.0% and 4.6 ± 3.5% of all LF waves, respectively.

**Figure 2.**
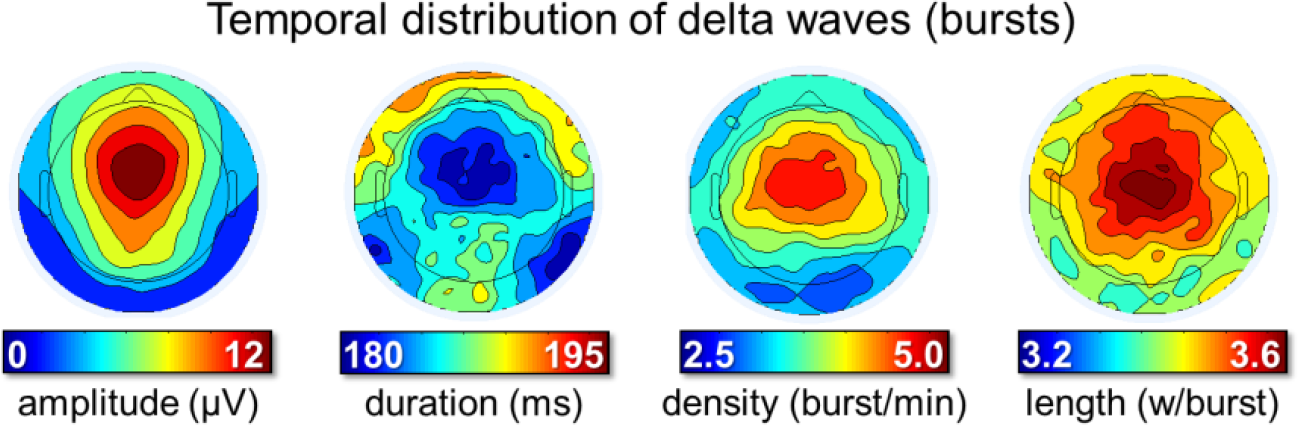
Temporal distribution LF waves (occurrence in bursts). Bursts were defined as series of at least 3 consecutive waves (1-5 Hz) whose maximum negative peaks were separated by less than 750 ms. From left to right, the plots display the density of LF wave bursts (number of bursts per minute), the mean amplitude (µV) and duration (ms) of waves included in a burst, and the number of waves included in the burst (expressed as waves per burst).

**Figure 3.**
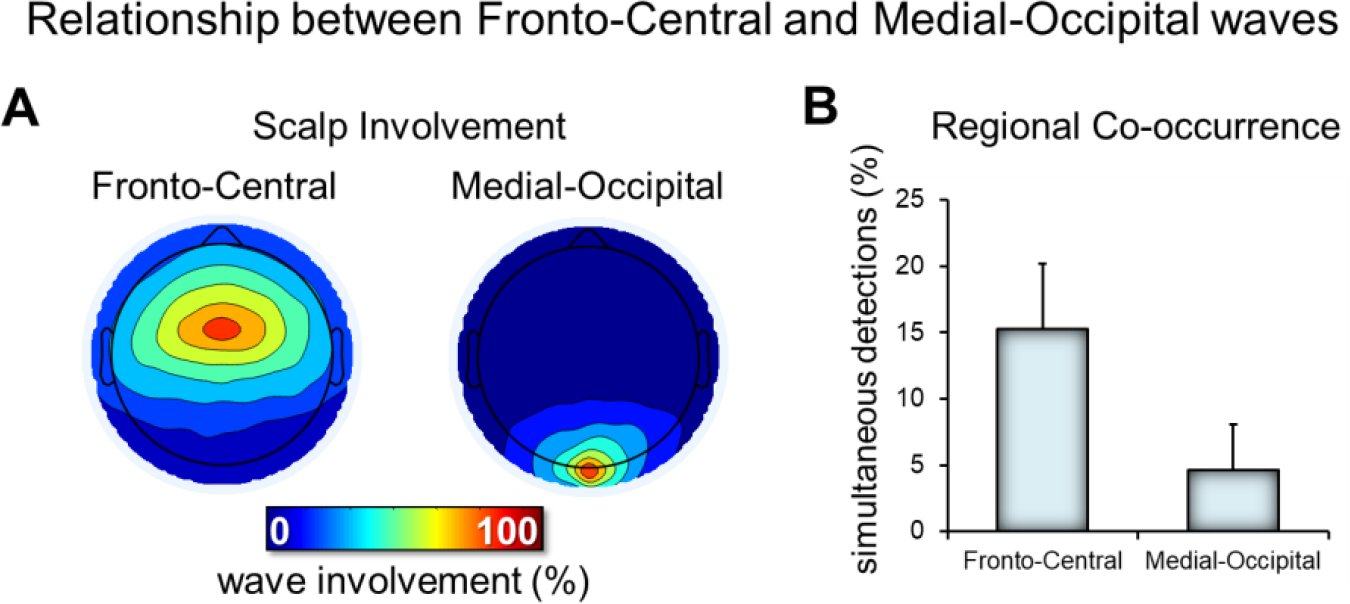
Relationship between fronto-central and medial occipital LF waves. Panel A shows the typical scalp involvement for the two types of LF waves. For each LF wave detected in the two electrodes of interest (fronto-central and medial-occipital), the number of concurrent detections (in a window of 200 ms, centered on the wave negative peak) in other electrodes was computed. Thus, each map shows the number of cases in which each electrode showed a co-occurring LF wave detection (the electrode of interest has 100% value in each map). Panel B shows the relative occurrence of simultaneous detections across the two electrodes of interest.

### Faster fronto-central waves (‘sawtooth waves’)

Visual inspection of these waves revealed that many of them looked like typical ‘*sawtooth waves*’ of REM-sleep (Figure 4A), displaying a notch either before or after the maximum negative peak. These waves showed a frequency peak between 2.5 and 3 Hz and a maximum power between the vertex (Cz) and frontal (Fz) electrodes (Figure 4A-B). Source modeling of the signal corresponding to the maximum negative peak revealed the greatest overlap between subjects (>10) in a medial frontal area comprising the mid-cingulate gyrus, the supplementary motor area (SMA) and the medial part of the primary motor (M1) cortex (Figure 4C). Sawtooth waves displayed a clear density increase before rapid EM and during phasic REM periods (Figure 4D), in line with previous observations in both human subjects and primates [32–37]. A direct positive correlation between the number of sawtooth waves preceding the EMs and the number of associated EMs was also observed (Figure 5). Finally, the analysis of *gamma* activity during sawtooth waves revealed a positive, in-phase, association between low- and high-frequency EEG activity (Figure 4E).

**Figure 4.**
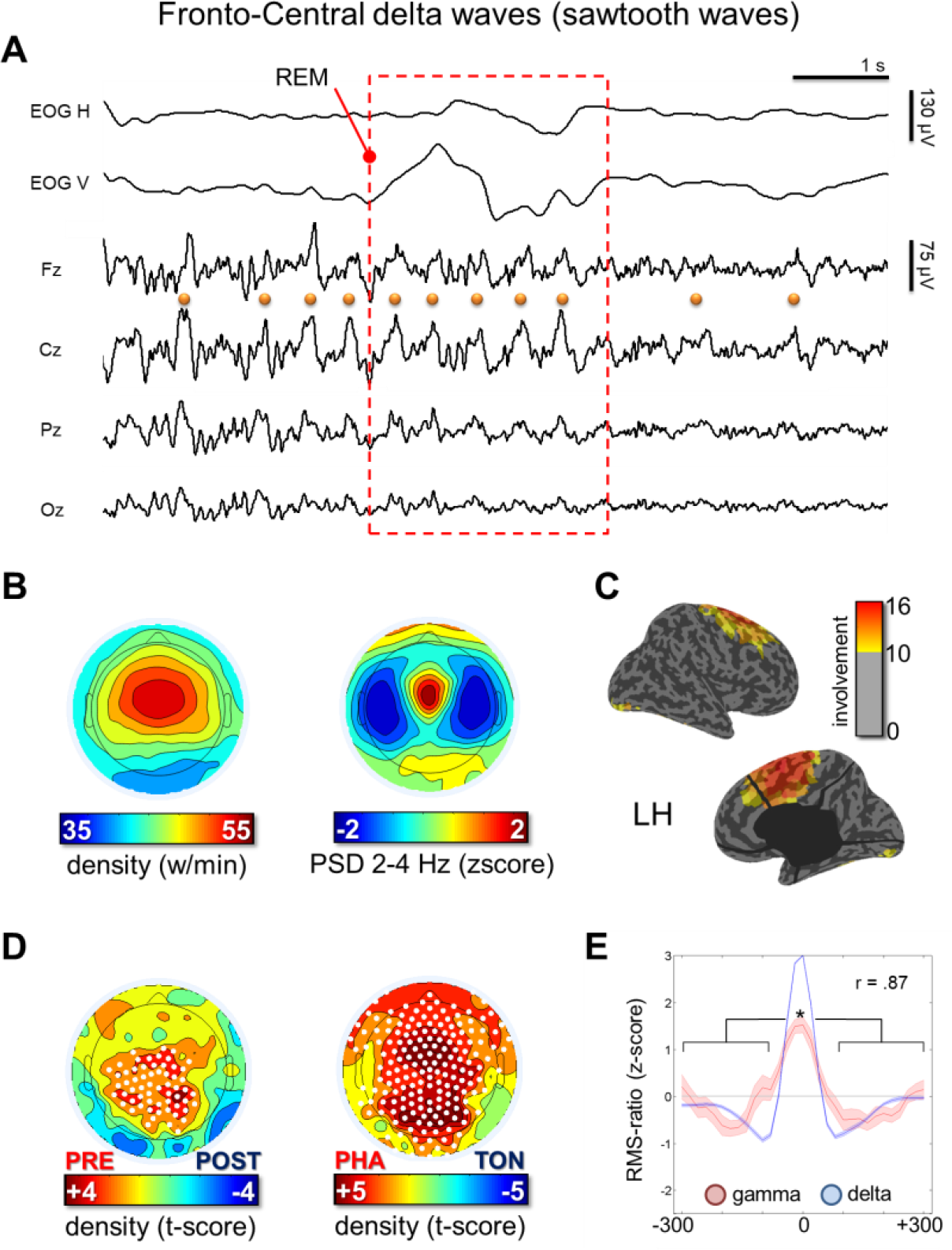
Panel A shows representative EEG traces containing fronto-central LF waves with a duration of 125-250 ms (half-wave). Most waves corresponded to typical notched sawtooth waves of REM-sleep. Panel B shows density- and power-based maps indicating their topographic distribution. Power spectral density (PSD) was calculated in 6s epochs and across all REM cycles using the Welch’s method (8 sections, 50% overlap) and integrated in the frequency-range of interest. Panel C shows the typical peak of activity of fronto-central waves in source space (overlap between subjects). Panel D shows a comparison between wave density before and after isolated eye movements and between phasic and tonic REM-periods. Panel E depicts the relationship between delta and gamma activity (RMS of band-limited signal). RMS traces were aligned using the maximum negative peak of each wave as a reference. * marks a significant increase of gamma activity at the wave peak with respect to baseline (p < 0.05).

**Figure 5.**
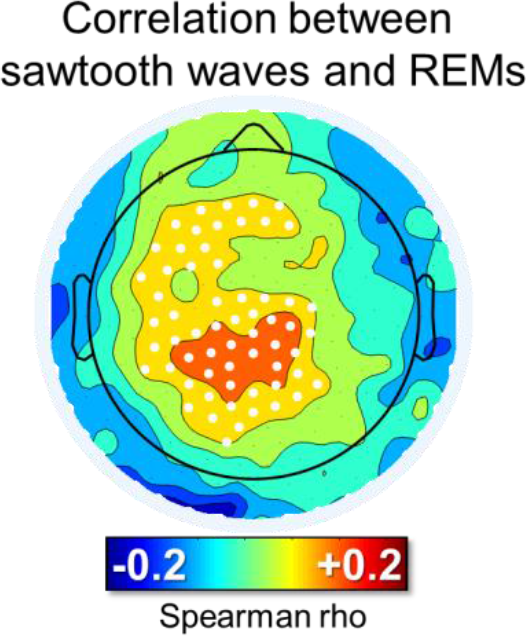
Correlation between the number of sawtooth waves preceding rapid eye movements and the number of subsequent rapid eye movements. For each isolated group of eye movements, the burst of the 125-250 ms half-waves (amplitude > 10 µV, inter-wave distance < 1 s) closest to the beginning of the first eye movement (maximum distance < 600 ms) was identified. White dots mark significant effects at group level (p < 0.05, corrected).

Next, given the notched appearance of typical sawtooth waves [38], we evaluated the topographic and cortical distribution of the signal during the notch and maximum negative peak of the wave separately. This analysis, performed at the scalp and source level revealed that the notched appearance results from the superimposition of two cortical sources: a fronto-central source corresponding to the maximal negative peak of the sawtooth wave, involving the mid-cingulate, medial and superior frontal areas (including motor and premotor regions), and a posterior source, corresponding to the notch of the wave, involving occipital and inferior temporo-occipital areas. Results of one representative subject are shown in Figure 6. For consistency, only sawtooth waves with a notch preceding the maximum negative peak were visually identified for this analysis although sometimes, a notch could also be observed following the maximum negative peak.

**Figure 6.**
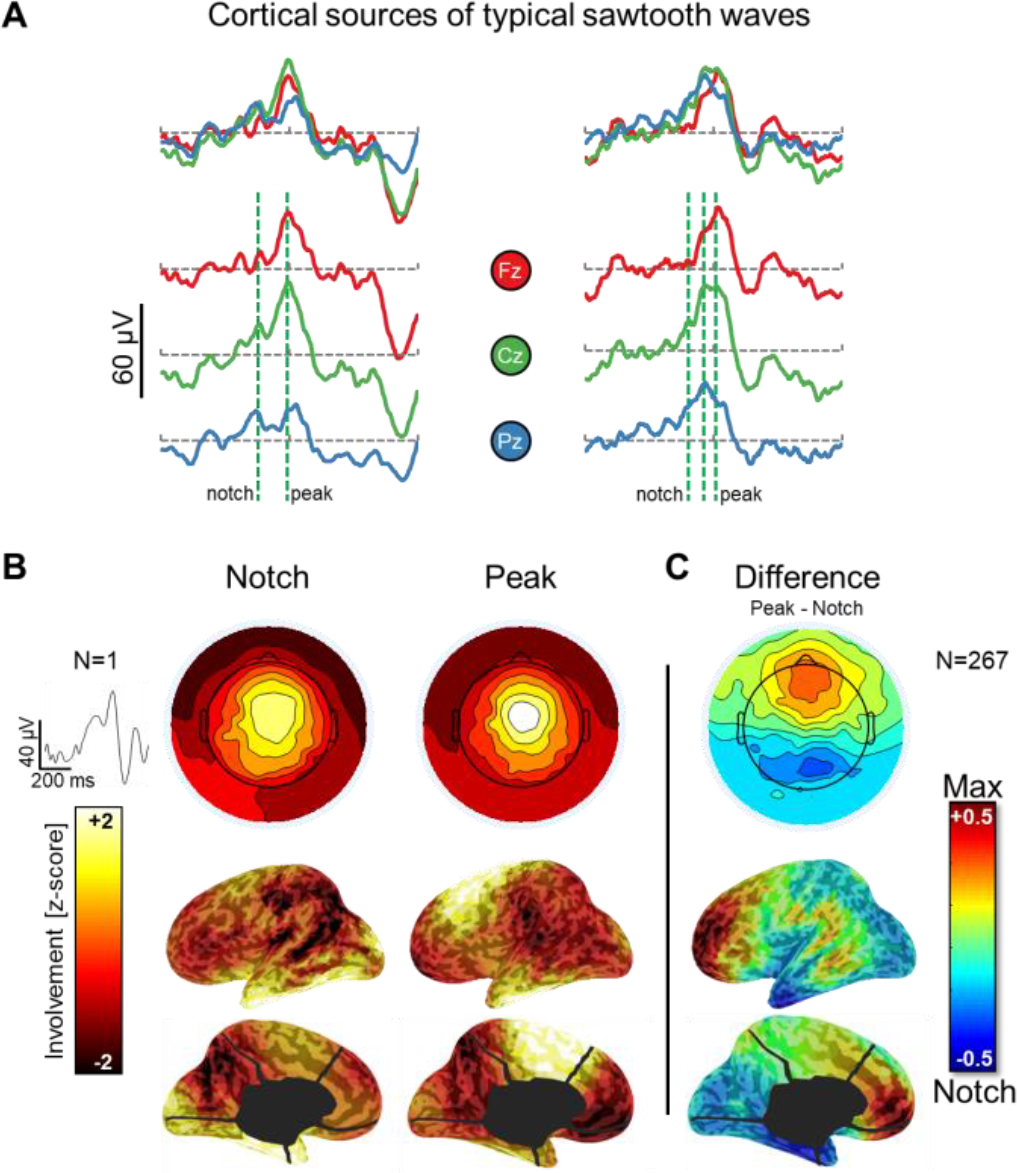
Cortical sources of typical sawtooth waves. Panel A shows the EEG traces (three electrodes, corresponding to Fz, Cz, and Pz) of two representative notched sawtooth waves (±250 ms around the negative peak of the algorithm detection). Panel B shows the scalp and cortical involvement corresponding to the notch and the maximal negative peak of a single sawtooth wave. Panel C shows the difference in average involvement between the notch and maximal negative peak (computed across 267 sawtooth waves in one representative subject, similar results were obtained in other participants). For this evaluation, all detections in one subject were visually inspected to identify typical sawtooth waves with a negative amplitude greater than 20 µV and a well-recognizable notched, triangular shape. The timing of the notch and the maximal negative peak were marked manually. Differences shown in Panel C are statistically significant over all voxels (p<0.05, corrected).

### Slow medial-occipital waves

Visual inspection of EEG traces revealed that the great majority of slow (300-500 ms) medial-occipital waves corresponded to shallow negative deflections, frequently accompanied by faster, superimposed theta-alpha activity (Figure 7A). These waves showed their maximum power in the occipital cortex (Figure 7A-C) and did not display clear changes in relation to rapid EM. However, they showed a relative decrease in centro-frontal, but not in occipital areas during phasic vs. tonic REM-sleep periods (Figure 7D). Similar to slow waves in NREM-sleep (e.g., [17]), REM slow waves were associated with a decrease in gamma activity (Figure 7E).

**Figure 7.**
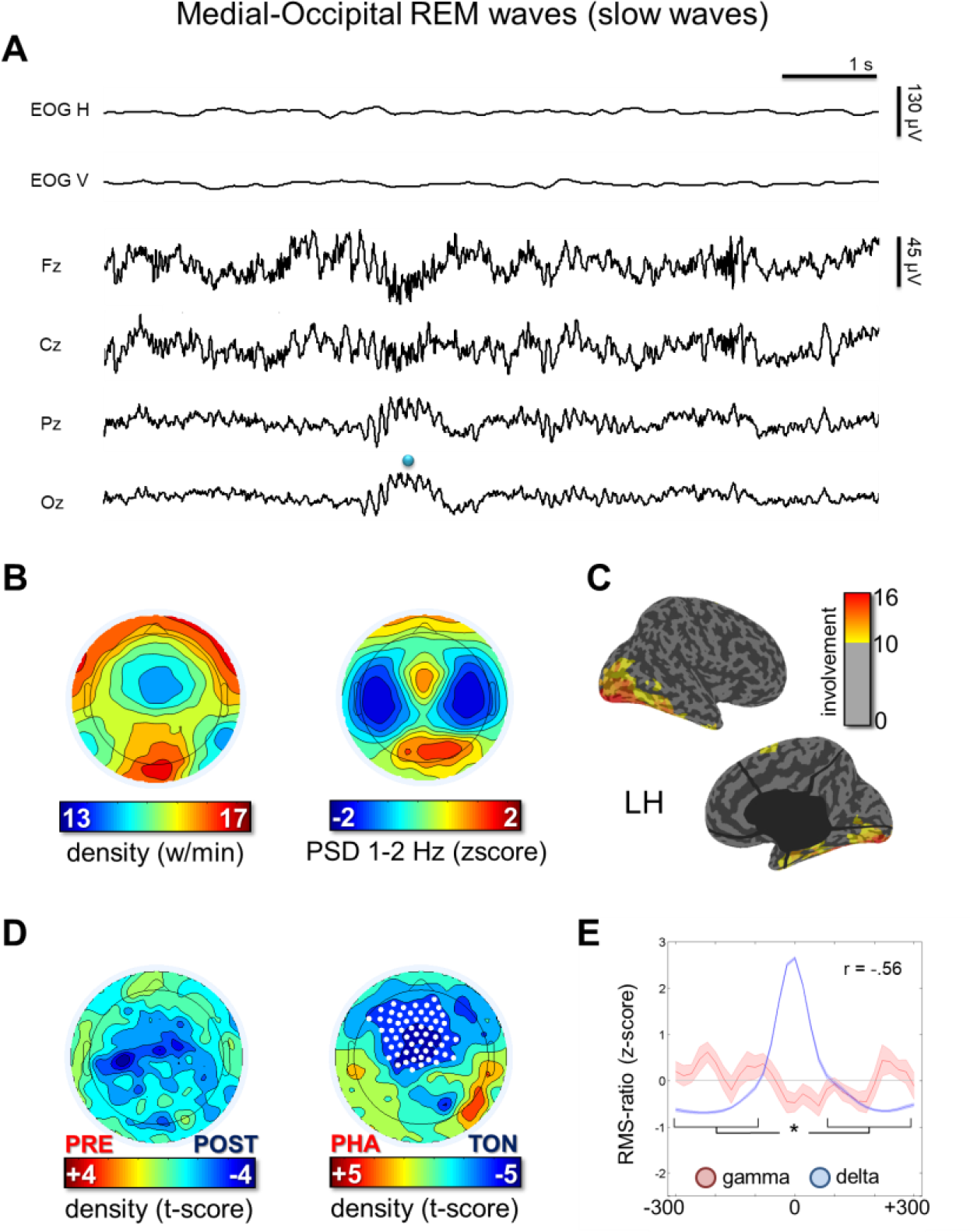
Panel A shows representative EEG traces containing medial-occipital LF waves with a duration of 300-500 ms (half-wave). Panel B shows density- and power-based maps indicating their topographic distribution. Power spectral density (PSD) was calculated in 6 s epochs and across all REM cycles using the Welch’s method (8 sections, 50% overlap) and integrated in the frequency-range of interest. Panel C shows the typical peak of activity of medial-occipital waves in source space (overlap between subjects). Panel D shows a comparison between medial-occipital wave density before and after isolated eye movements and between phasic and tonic REM-periods. Finally, panel E depicts the relationship between delta and gamma activity (RMS of band-limited signal). RMS traces were aligned using the maximum negative peak of each wave as a reference. * marks a significant increase of gamma activity at the medial-occipital wave peak with respect to baseline (p < 0.05).

### Overnight changes in LF waves

An analysis of overnight changes in wave amplitude revealed a significant, non-region-specific (i.e., both frontal and occipital) and non-frequency-specific (i.e., for both faster and slower waves) decrease across the night (Figure 8). No overnight changes in wave density were observed.

**Figure 8.**
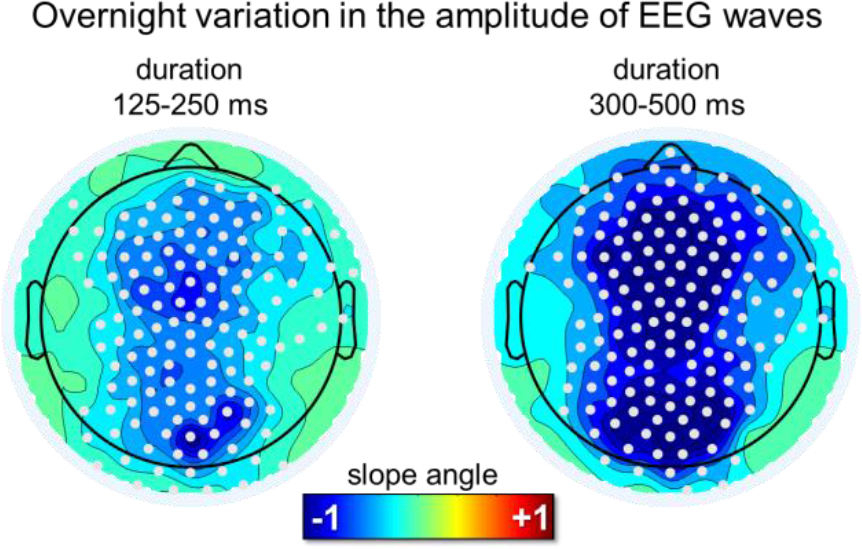
Overnight changes in the amplitude of LF waves in REM sleep for the ‘sawtooth’ frequency range (125-250 ms) and the medial-occipital wave range (300-500 ms). A diffuse decrease in amplitude was observed for both frequency ranges. White dots mark significant effects at group level (p < 0.05, corrected).

### Relationship with low-frequency oscillations of NREM sleep

We then asked whether waves with ‘sawtooth’ and ‘medial-occipital’ characteristics were specific to REM-sleep or whether they could also occur in NREM-sleep. We found that the density of LF waves in the sawtooth frequency range differed significantly between stages (Friedman test p < 0.0001, X^2^= 20.0) in the centro-frontal region, being maximal in REM-sleep, and lowest in N3-sleep (p < 0.005, |z| = 2.803; Wilcoxon signed-rank test). Instead, the density of slower waves in the medial-occipital cluster did not change significantly across sleep stages (Friedman test p > 0.05, X^2^= 5.4; Figure 9). Amplitude was significantly higher in N2 and N3 sleep than in REM-sleep for both faster fronto-central waves (Friedman test p < 0.0001, X^2^= 16.8.0) and slower medial-occipital waves (Friedman test p < 0.0001, X^2^= 20.0). For a topographic display of waves across stages, see Figure 10.

**Figure 9.**
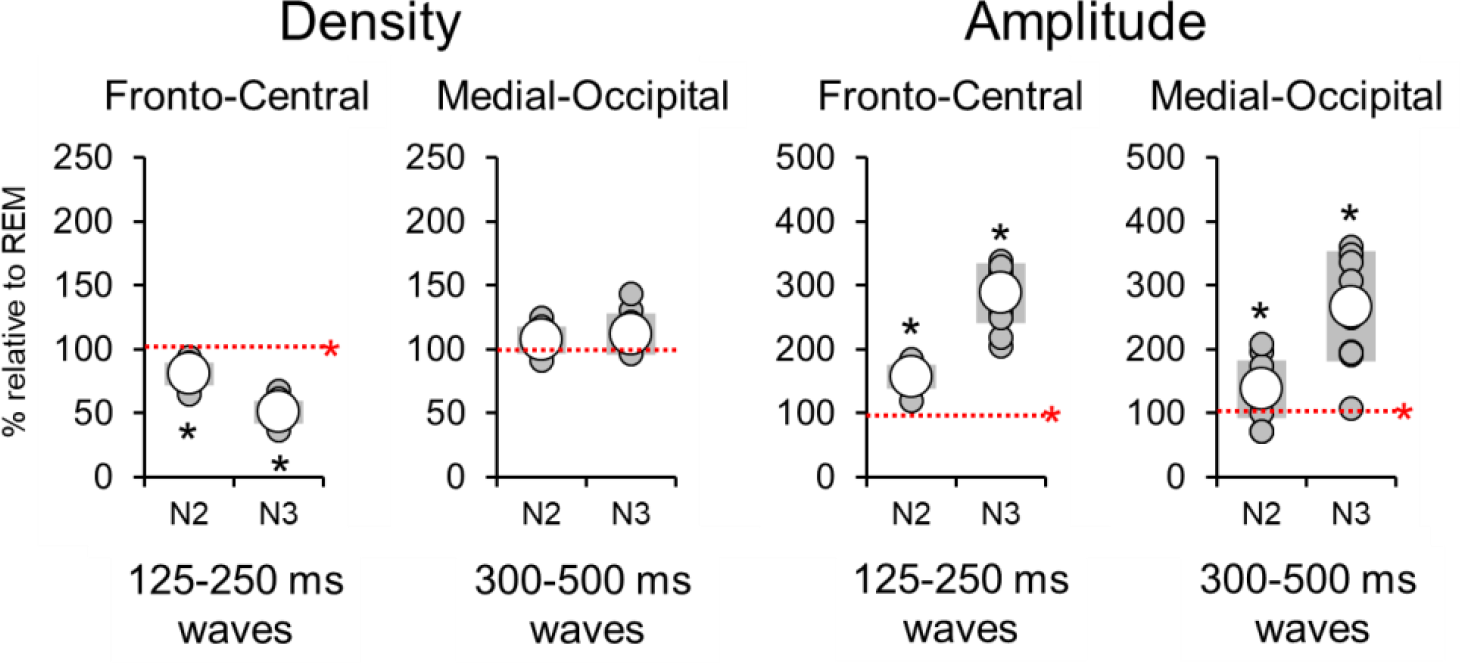
Relative variations in the density and amplitude of fronto-central and medial-occipital LF waves in N2 and N3 with respect to REM sleep. The dashed red line corresponds to the level observed in REM sleep (100%). * denotes significant effects at p < 0.05 (red for Friedman test for stage-effect, black for post-hoc Wilcoxon signed rank tests)

**Figure 10.**
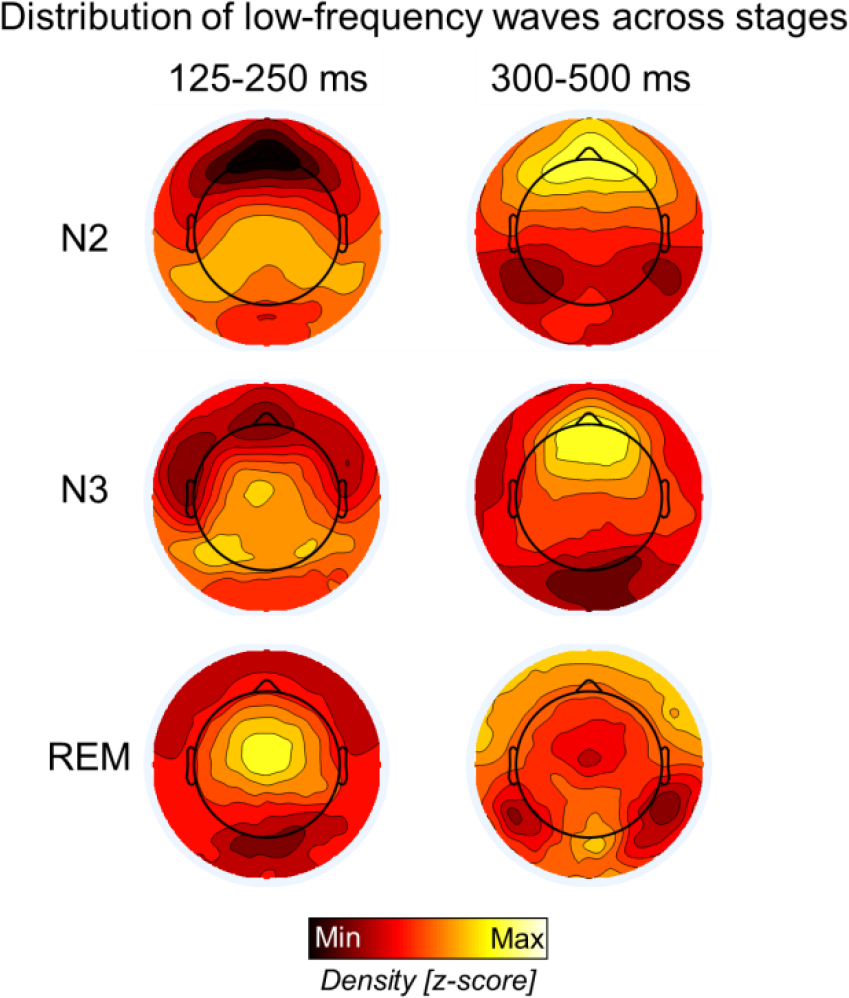
Relative topographic distribution of EEG waves with durations of 125-250 ms (left) and 300-500 ms (right) in N2, N3 and REM sleep. Density values were z-scored to facilitate comparison across stages. The strong frontal slow wave activity in NREM sleep likely masks the smaller low-frequency oscillations in posterior regions.

### Relationship with low-frequency oscillations of wakefulness

Because we found that the slower medial occipital waves were not specific to REM-sleep, but also occurred with a similar incidence in NREM-sleep, we asked whether they were also present in wakefulness. To this aim we performed an additional analysis in a different dataset, containing both REM-sleep and wake data (2 min of relaxed wakefulness obtained with eyes closed in the morning). This analysis showed that medial-occipital slow waves had a significantly lower density (p < 0.005, |z| = 2.934) and amplitude (p < 0.005, |z| = 2.934) in wakefulness relative to REM-sleep. Specifically, the average density was 5.5 ± 2.8 w/min in awake subjects, and 13.9 ± 1.7 w/min during REM-sleep (across the whole night), suggesting that they are clearly modulated by the change in behavioral state (sleep vs. wakefulness).

## Discussion

Here we show that LF waves (≤4 Hz) are an integral part of human REM-sleep. By analyzing LF wave properties separately, two distinct spatial clusters spontaneously emerged: a frontal-central cluster with frequent, faster (2-4 Hz) high-amplitude waves and a medio-occipital cluster with isolated, slower (<2 Hz), low-amplitude waves. Both oscillations were essentially local, involving only a minority of channels, and occurred independently from one another.

### Fronto-central sawtooth waves

These waves resembled typical sawtooth waves of REM sleep, originally described in the 1960s [32,39,40]. There is no universal agreement on defining criteria for sawtooth waves, but they are often described as medium amplitude waves (≥ 20 µV) between 2 and 5 Hz [32,39,41–43], with a triangular shape (initial slow increase followed by a steep decrease [38,44,45]), and a maximal fronto-central amplitude [41,46]. The presence of a notch, usually on the positive-to-negative slope, is occasionally mentioned [38,47]. Sawtooth waves tend to occur in bursts [35,48], herald the beginning of REM-sleep [35] and often, but not necessarily accompany bursts of rapid EMs [32,33,35,37]. The characteristics of fronto-central waves described herein are consistent with these observations. In addition, we provide several novel findings. First, using source modeling, we demonstrate a maximal involvement in medial prefrontal areas (mid-cingulate cortex, SMA and the medial M1). We also identified a secondary temporo-occipital source, responsible for the sawtooth wave ‘notch’. Second, we showed that sawtooth waves are accompanied by an (in-phase) increase in high-frequency activity, suggesting an ‘activating’ effect (i.e., increased neuronal firing [49]). The mechanism underlying these waves is thus likely different from NREM slow waves, which are associated with neuronal silence (*OFF-periods*). Finally, we were able to document a direct correlation between sawtooth waves and rapid EMs (the more sawtooth waves, the more subsequent EMs).

The characteristics of sawtooth waves raise the possibility that they are related to ponto-geniculo-occipital (PGO) waves, which were originally described in cats and consist of phasic electric potentials recorded from the pons, the lateral geniculate body (LGB), and the occipital cortex [50–53]. They have also been mapped in limbic areas (amygdala, cingulate and hippocampus). Similar to sawtooth waves, PGO waves last between 60 and 360 ms [54], appear at the NREM-REM sleep transition, several seconds before the first rapid EM [35], occur in bursts (but isolated spikes are also observed [55]), and are typically, but not always associated with trains of EMs [56]. Although PGO waves have been recorded in many species, including nonhuman primates, their existence in humans remains controversial. A relation between sawtooth and PGO waves has been suggested based on the similar temporal characteristics, including their association with rapid EMs [37,57], but the different cortical distributions (fronto-central vs occipital) have hampered further analogies. The temporo-occipital component that we identified here may actually represent the cortical manifestation of human PGO waves. The increases in gamma power associated with sawtooth waves fit well with this hypothesis. Indeed, PGO waves have been named ‘cortical activation waves’ [58] because they are associated with increased neuronal firing in several cortical regions. Consistent with our findings, gamma power increases have been reported during rapid EM in frontal and central regions in REM sleep [59,60] and during phasic compared to tonic REM sleep [61]. Among the few case studies documenting potential human PGO-wave equivalents using intracranial recordings [62–64], one described the presence of simultaneous sawtooth waves in the scalp EEG, in addition to increases in gamma activity (15-35 Hz), supporting the analogy between sawtooth and PGO waves [64].

The correlation between sawtooth waves and rapid EM raises the question of whether sawtooth waves are directly involved in rapid EM generation. However, several points argue against this possibility. First, although saccades during wakefulness and REM sleep share many common features, including analogous ERP waveforms, the same phase reset of theta activity in medial temporal regions, a comparable modulation of neuronal firing rates [65] and the activation of similar brain regions [66,67], sawtooth waves do not accompany rapid EM during wakefulness. Second, while sawtooth waves clearly encompass the frontal and supplementary eye field, they also extend beyond these regions, to V1, the SMA and limbic regions (midcingulate inferior/middle temporal cortex). Activations of these areas [59,67–70] and the LGB [71-73] have been shown to differentiate brain activity during REM sleep from EM during wakefulness [71,73]. Our results suggest that these differences may be mediated by sawtooth waves, which are concomitant to EMs. Finally, the fronto-central topography is almost identical to some NREM potentials that are not associated with EMs, including vertex waves [17,41] and high-frequency activity increases following type I slow waves [17–19]. During wakefulness, vertex potentials can be induced by sudden and intense stimuli of various sensory modalities, including pain. These potentials have been suggested to be related to stimulus saliency, that is, the ability of a stimulus to stand out from its surroundings [74]. Rapid EMs following sawtooth waves may thus constitute a reaction to a salient, possibly dreamt event. Consistent with this, PGO spikes can be induced in cats using sounds, during both quiet wakefulness, REM and NREM sleep, and may accompany a ‘startle-like’ reaction [75], suggesting common behavioral significance of these potentials across states.

### Medio-occipital slow waves

The second cluster of LF oscillations consisted of occipital slow waves (<2 Hz). This finding is consistent with previous work, describing the maximal peak of low-delta activity in occipital areas during REM-sleep (e.g., [14,15]). Our results demonstrate that this source of delta activity is due to small, local slow waves that are not modulated by phasic REM and are associated with a suppression of gamma activity, similar to NREM slow waves (e.g., [17]). We also show, for the first time, that occipital slow waves occur with a similar density frequency, but different amplitudes, across all sleep stages. The amplitude of LF waves was higher in N3 relative to N2 or REM sleep and decreased across subsequent REM-sleep periods. These variations may reflect neuromodulatory and sleep-dependent changes in synaptic density/strength which may affect the efficiency of neural synchronization during the generation and spreading of slow waves [24,76]. On the other hand, occipital slow wave density remained similar in NREM and REM sleep, but was higher than in wakefulness. Consistent with this, the occipital cortex, relative to other cortical areas, shows the smallest changes in SWA at NREM-REM sleep transitions [15,77]. The relatively low frequency of occipital REM slow waves (<2 Hz) is in line with the recent observation of a lower frequency of occipital NREM slow waves (∼1 Hz) relative to anterior areas [78].

Our observation of local occipital slow waves occurring at a similar rate across NREM and REM-sleep is consistent with the hypothesis that slow waves in primary cortices could be responsible for sensory disconnection during sleep [13]. However, with respect to slow waves recently described in mice [13], several differences should be noted. First, we did not observe SWA peaks in primary cortices other than V1. It is possible that local slow waves in other primary cortices may have been masked by the preponderant sawtooth activity in these areas. Indeed, in a recent study comparing SWA between REM and wakefulness, SWA activity was found to be higher in primary sensory cortices in REM sleep after excluding sawtooth waves [79]. This possibility is also in part supported by the observation of a relative decrease in the fronto-central wave density in the 1-2 Hz range during REM periods dominated by ‘activating’ sawtooth waves (i.e., phasic REM). Second, in our study we found that occipital slow waves, as opposed to sawtooth waves, were not modulated by phasic REM-periods like in the animal study. This may be due to the different definitions of phasic REM periods that were used, based on rapid EMs in humans and on whisking events (vibrissal EMG) in mice. However, we cannot exclude that slow waves observed in these two studies may represent distinct phenomena.

### Limitations

We focused on a minority of LF waves that were typical of clusters emerging from topographical analyses. Thus, there is a large ‘unclassified’ background of LF waves that remains unexplored in the present study. Due to volume conduction affecting scalp EEG recordings, masking effects of prominent sawtooth waves with respect to other slow waves could not be avoided. Future human studies using intracranial recordings should explore the regional distribution of slow waves in REM sleep (primary vs. secondary sensory cortices) and their association with *off-periods*.

### Conclusions

LF oscillations (≤4 Hz) are an integral feature of REM-sleep. There appear to be at least two distinctive clusters of LF oscillations with different properties: a fronto-central cluster characterized by faster, activating ‘sawtooth waves’ that share many characteristics with PGO waves described in animals, and a medio-occipital cluster containing slow waves which are more similar to NREM sleep slow waves.

## Acknowledgments

The authors thank Brady Riedner, Michele Bellesi, Josh J. LaRoque and Xiaoqian Yu for technical assistance and help with data collection.

